# Toxicological Analysis of Poly-Lactide Co-Glycolide Microparticles, Chitosan Nanoparticles and Poly(Anhydride) Nanoparticles in Vero Cell

**DOI:** 10.1101/2022.03.17.484425

**Authors:** Dipankar Hazarika, Durlabh Prasad Borah, Jayanta Sarma Boruah, Yasmin Begum, Anisha Sultana, Shantanu Tamuly, Rita Nath, Devasish Chowdhury, Biswajit Dutta, Probodh Borah, Dhruba Jyoti Kalita

## Abstract

The present study was carried out to evaluate *in-vitro* toxicity associated with chitosan nanoparticles, poly(anhydride) nanoparticles and poly-lactide co-glycolide (PLG) microparticle in Vero cell line. The cytotoxicity of all three micro/nano-particles was assessed using different concentration. For each concentration, the confluent monolayer was treated for a period of 36-48 hours and studied for morphological alteration, live and death count etc. after the treatment. It was observed that the different concentrations of chitosan nanoparticles and Gantrez^®^ nanoparticles did not have significant effect on the cell viability as evident from the non-significant difference between the OD_540_ of formazan product formed from MTT in treated and untreated cells. The concentration of chitosan nanoparticles and Gantrez^®^ nanoparticles up to 1000 μg/ml did not have any influence in cellular metabolic activities and viability. However, a reduction in the cellular viability and metabolic activities were observed when PLG microparticles were used at 1000 μg/ml. At concentrations below 1000 μg/ml, all the nanoparticles/microparticles were found to be safe in terms of cytotoxicity.

---

The advantage of oral administration of vaccines in poultry is obvious as compared to the parenteral route. First, it is faster and much easier to administer for mass application without requiring highly trained personnel and carries no risk of needle stick injury or cross-contamination (McCluskie *et al*, 2000). Nanoparticles and microparticles offer great applications in the field of biological sciences in terms of oral drug and vaccine delivery systems. Nanoparticles made from natural/synthetic polymers, lipids, proteins and phospholipids have received greater attention due to higher stability and the opportunity for further surface modifications (Stark, 2011). They can be tailored to achieve both controlled drug release and disease-specific localization, either by tuning the material characteristics or by altering the surface chemistry (Herrero-Vanrell *et al*., 2005). *In vitro* assays form a battery of first-line methods for discriminating between safe and hazardous nanoparticles. Such assays may also provide insight into specific mechanistic pathways such as potential to generate oxidative stress or inflammatory cytokines as well as cellular internalization of nanoparticles (Patravale *et al*., 2012). Since scanty of information are available on *in vivo* and *in vitro* toxicological evaluation of PLG microparticles, chitosan and Gantrez^®^ nanoparticles (poly(anhydride) nanoparticles) as oral vaccine carrier. Hence, the present study is undertaken to evaluate the *in vitro* assessment of cytotoxic effect of oral vaccine delivery system (poly-lactide co-glycolide microparticles, chitosan and Gantrez^®^ nanoparticles) in suitable cell line.

## MATERIALS AND METHODS

The present study was carried out in the laboratories of Department of Veterinary Biochemistry, and Veterinary Microbiology, College of Veterinary Science, Assam Agricultural University, Khanapara, Guwahati-781022. The chitosan nanoparticles, PLG microparticles and Gantrez^®^ nanoparticles were prepared as per the method described by Chowdhury et al. (2012) Zhao and Rodgers, (2006) and Camacho *et al*. (2011) respectively. The chitosan nanoparticles, Gantrez^®^ nanoparticles and PLG microparticles were characterized in terms of zeta size and zeta potential by dynamic light scattering (DLS) using Malvern Zeta-sizer Nano series. The cytotoxicity of all three micro/nanoparticles were assessed in Vero cell line using three different concentration (10 μg/ml, 100 μg/ml and 1000 μg/ml). For each concentration, the confluent monolayer was treated for a period of 36-48 hours and studied. At every 24 h interval, cells were observed for visible morphological changes under inverted microscope (Karl-Zeiss, Germany) and microparticle/nanoparticle treated cells were compared with the untreated cells. After 72h, the cells were trypsinized and the live and dead cell counts were determined using 0.4% (w/v) trypan blue exclusion method using hemocytometer and the MTT assay was done as per the method described by Bahuguna *et al*. (2017).

All the values were expressed as mean ± standard error (SE). The normality of the data was determined by Shapiro Wilk test. The between group difference of mean was determined by one way ANOVA (normally distributed data) or Krushkal-Walis test (non-normally distributed data). All of the statistical analysis was carried out in statistical software R (R Core Team, 2021).

## RESULTS AND DISCUSSION

The zeta size for chitosan nanoparticles, Gantrez^®^ nanoparticles and PLG microparticles were found to be 247.4, 262.8 and 3654 nm (Fig 1.). The cytotoxicity of all three micro/nano-particles was assessed in Vero cell line using different concentration. For each concentration, the confluent monolayer was treated for a period of 36-48 hours and studied for morphological alteration, live and death count etc. after the treatment.

**Fig. 1.**
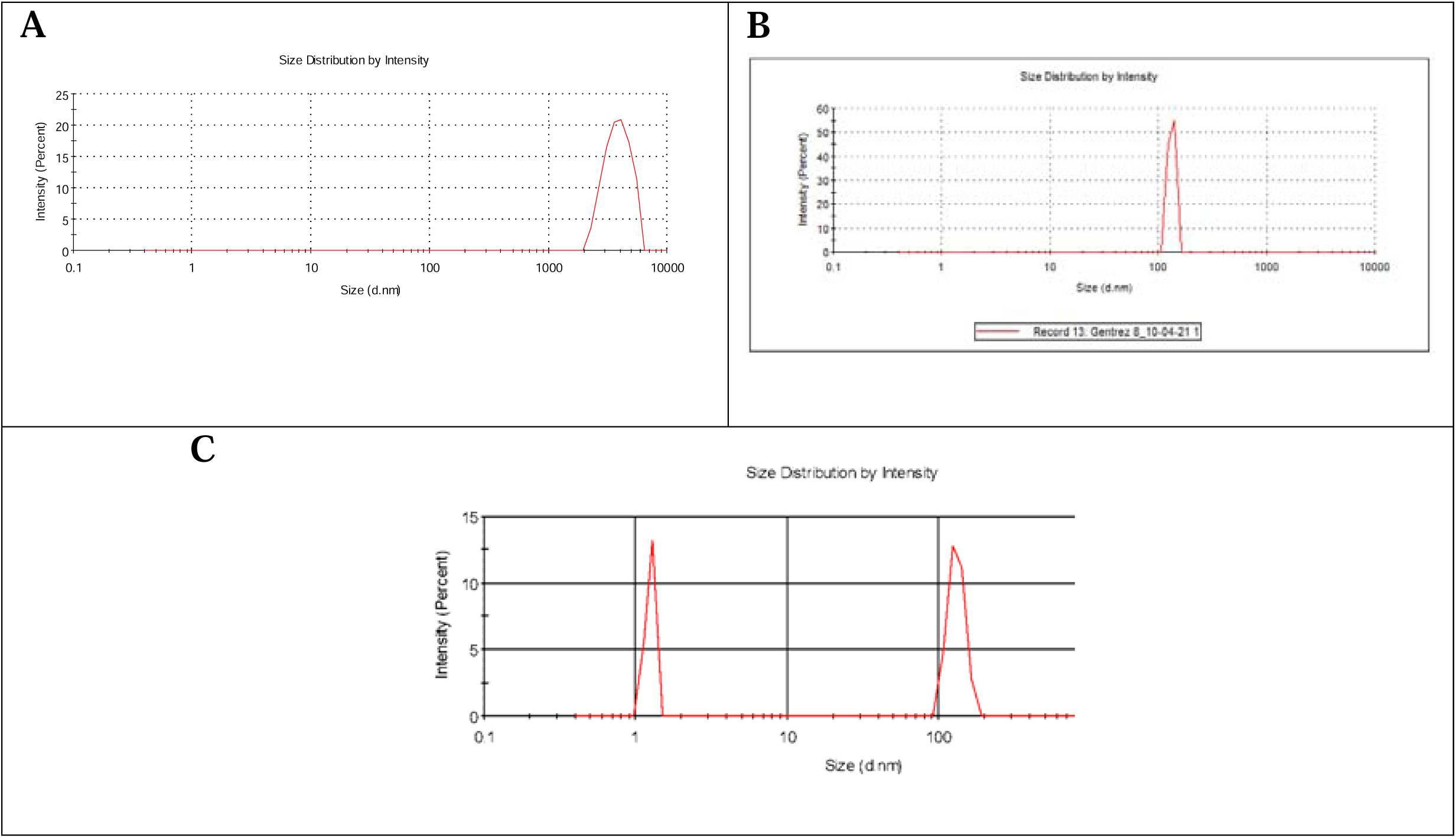
Zeta size of PLG microparticles, chitosan nanoparticles and poly(anhydride) nanoparticles.

### Morphological evaluation of treated cells

Morphological evaluation of the treated cells of Vero cell-line was done by comparing the cells with the control untreated flask. All the cells treated with different concentrations of PLG microparticles or Gantrez^®^ nanoparticles or chitosan nanoparticles were found to maintain the cell monolayer intact with minimal to no loss of cellular integrity and architecture like that of untreated cells (Fig. 2). The chitosan is degraded in biological systems under the influence of lysozyme and family of enzymes known as chitinases (Kean and Thanou, 2010). The chitosan nanoparticles in the range of 200 to 300 nm have shown to induce some toxicity in the zebra fish model by (Gao et al., 2011).

**Fig. 2.**
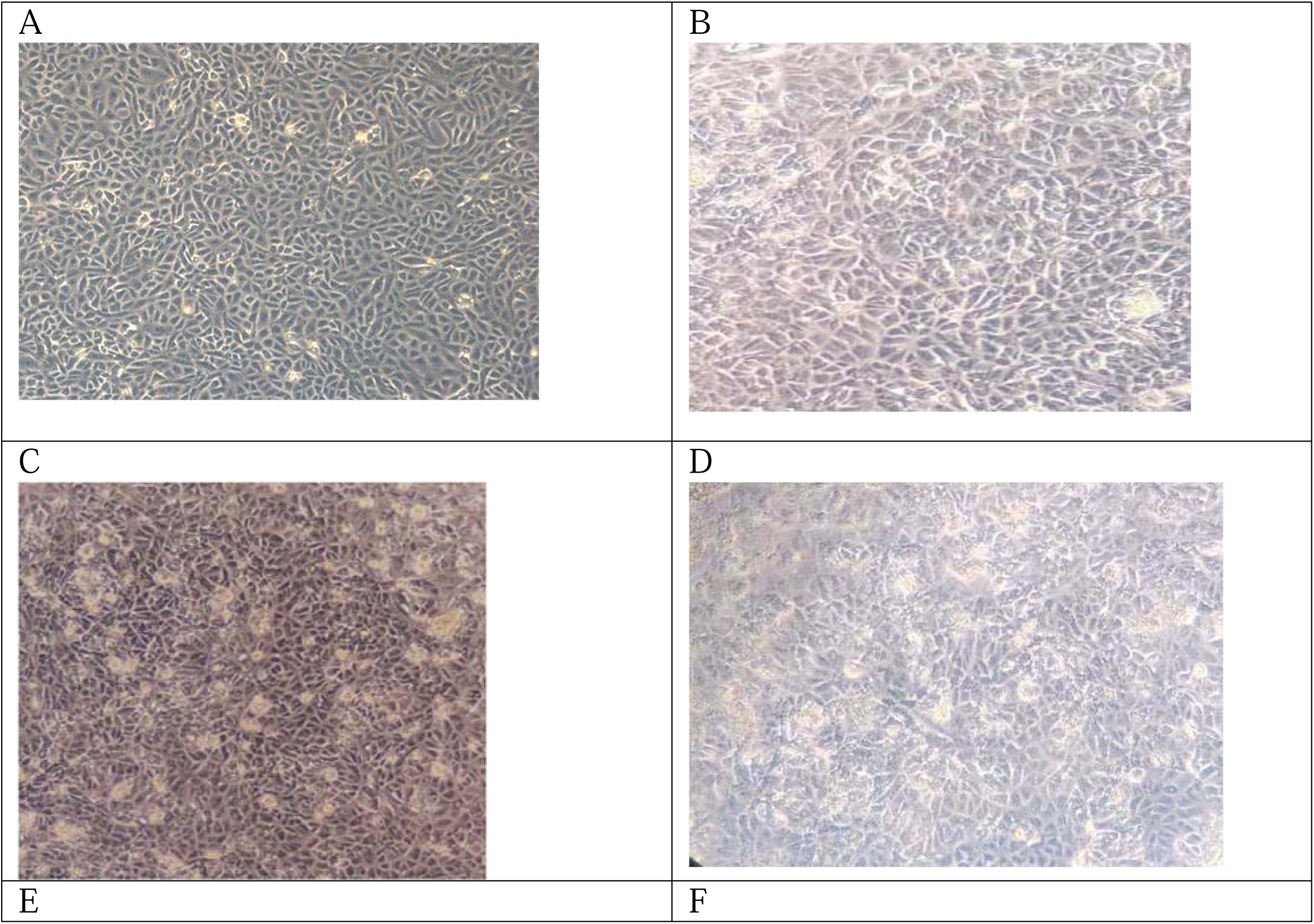
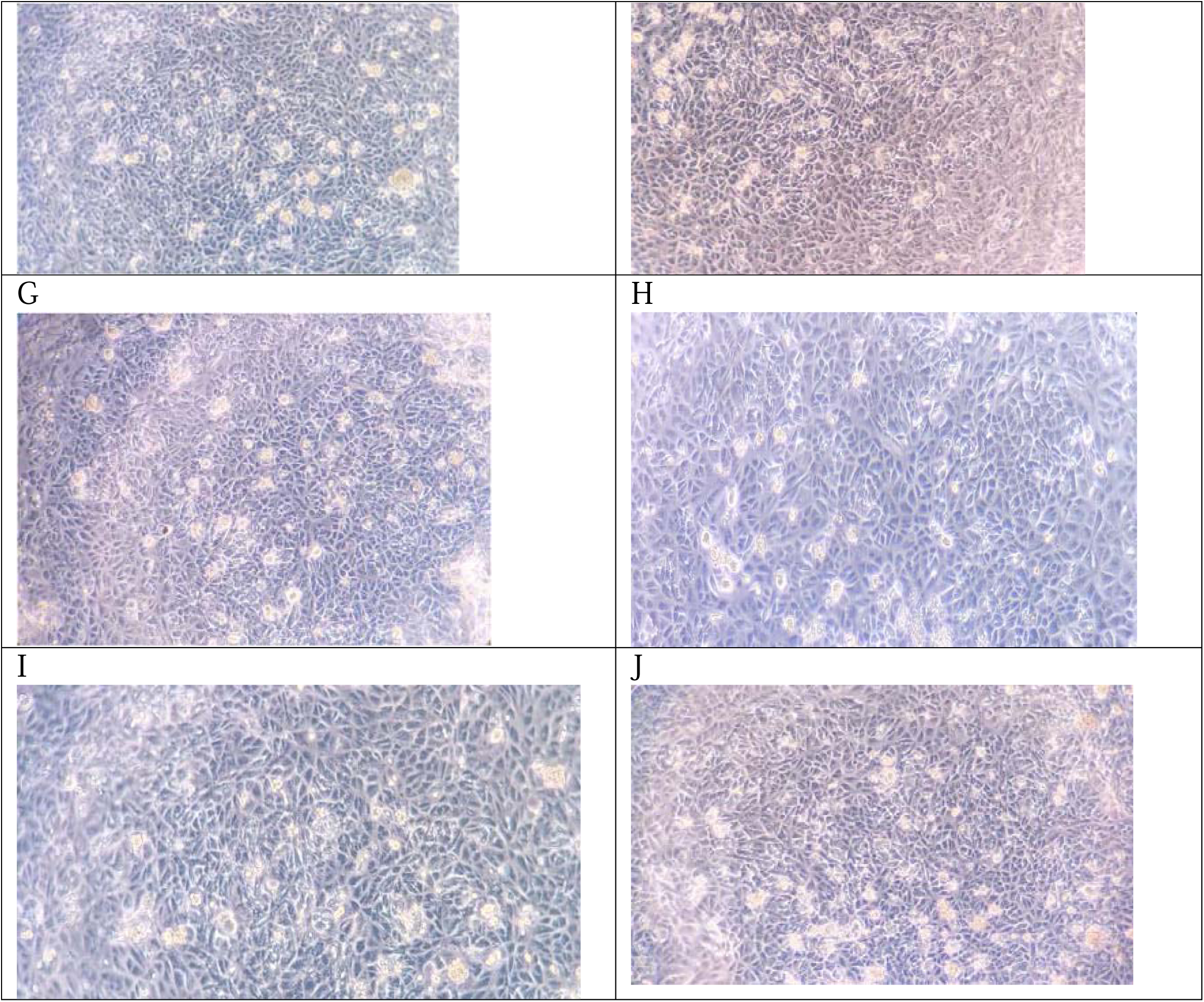
Morphological examination of A. untreated vero cell line, B, C, D: chitosan nanoparticle treated cell lines (10, 100, 1000 μg/ml), E,F,G: Gantrez® nanoparticle ((10, 100, 1000 μg/ml), H, I, J: PLG microparticles.

### Cell Viability

The cell death was determined by rounding, granulations, loss of intra-cellular matrix, clumping and finally detachment of the cells from the surface were the major manifestations. Results revealed that all three nanoparticles and their respective concentrations used were not cytotoxic to the Vero cells. The live and dead cell count was at par with the untreated control flask. However, there was slight reduction in the live cell count in treated flask as compared to untreated control flask but it was statistically non-significant. In the control flask, the live cell count was estimated to be 2.3 × 10^4^ cells / ml, while in the flasks treated with nano particles, it was ranging from 1.8-2.06 × 10^4^ cells / ml. There was no significant influence of different concentrations of nanoparticles and microparticles (Fig. 3). In addition, it was observed that the different concentrations of chitosan nanoparticles, Gantrez^®^ nanoparticles and PLG microparticles did not have significant effect on the cellular metabolic activity as evident from the non-significant difference between the OD_540_ of formazan product formed from MTT in treated and untreated cells (Fig 4).

**Fig. 3.**
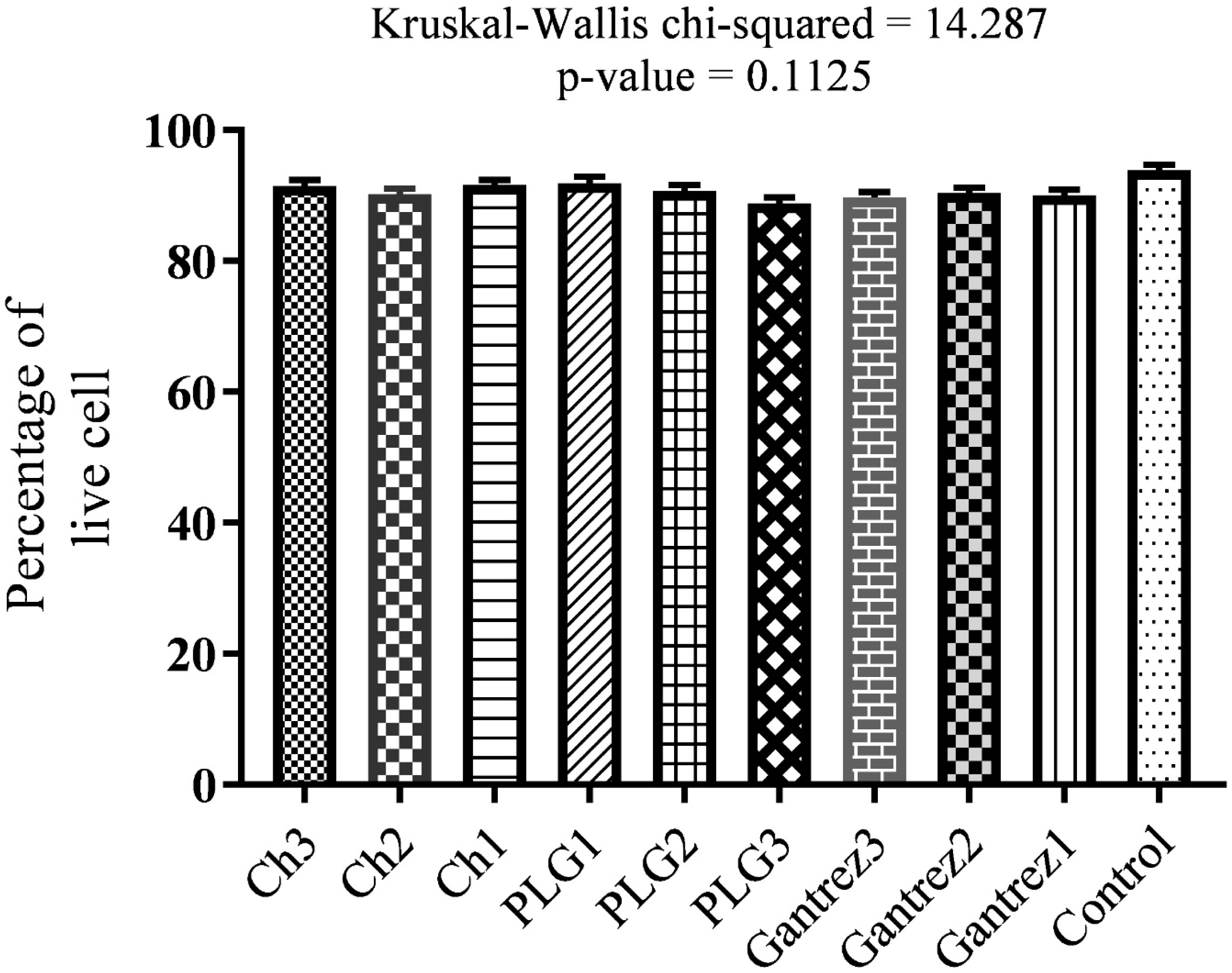
Cell viability of Vero cell lines treated with different nano-microparticles.

**Fig. 4.**
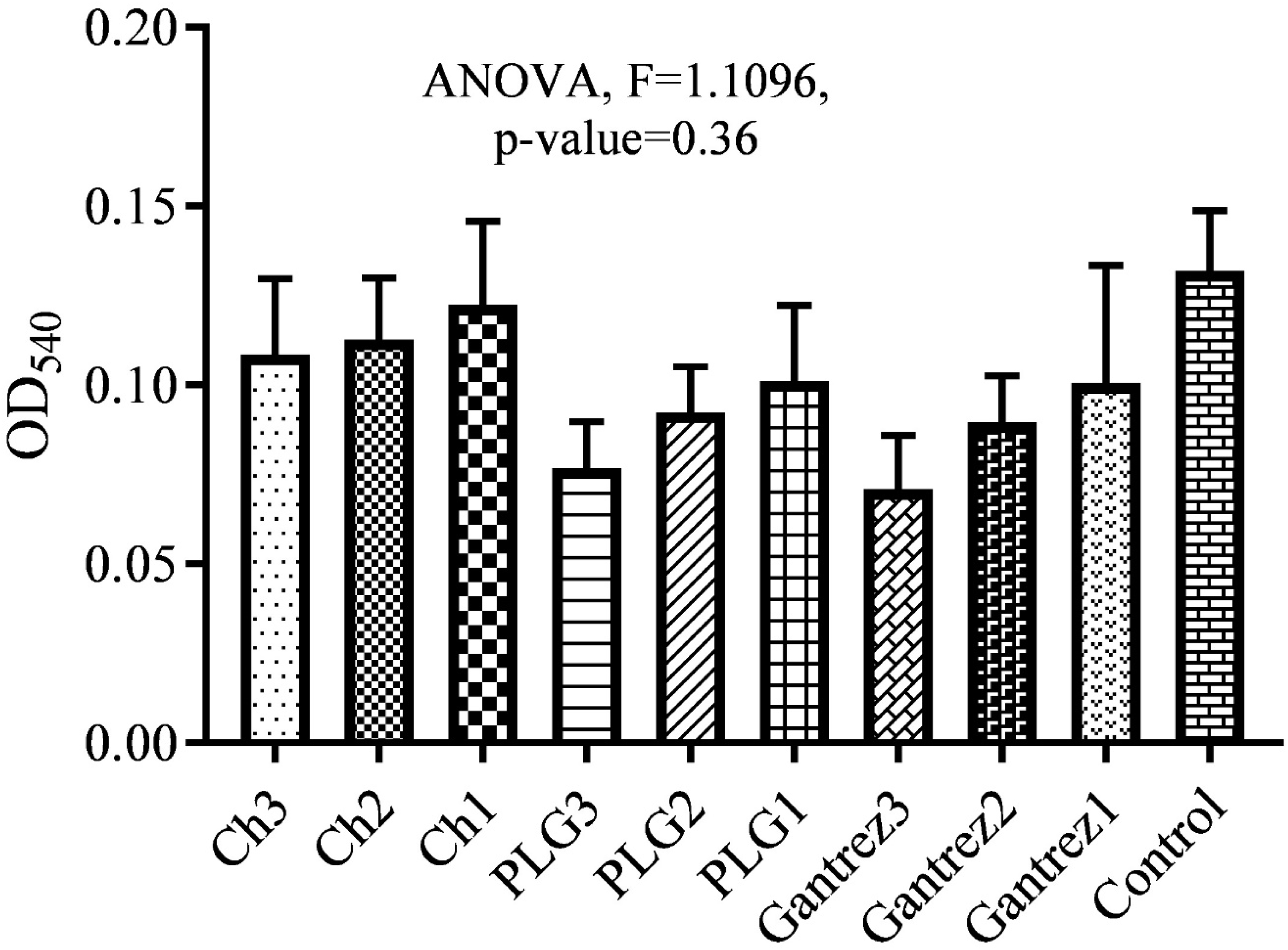
MTT assay of Vero cell treated with different concentration nano/microparticles.

The non-cytotoxic effects of the all concentrations of chitosan, Gantrez^®^ nanoparticles and PLG microparticles might be due to the biocompatibility and nontoxic nature of the particles which does not exert toxic effect on Vero cell. Similar observations were also reported by Ojer *et al*. (2013) while evaluating the cytotoxic effect of poly (anhydride) nanoparticles in HepG2 and Caco-2 cells using different concentrations and incubation time (0.0625, 0.125, 0.25, 0.5, 1 and 2 mg/ ml for 4, 24, 48 and 72 h) and the result showed that the viability starts to decrease only at very high concentrations (1 and 2 mg/ ml) and long incubation times (48 h). Further, it has been reported that cytotoxicity effects of chitosan nanoparticle (CSNPs) is relatively low, concentration-dependent and influenced by particle size. CSNPs were relatively nontoxic regardless of particle size at low CSNPs concentrations (10 and 100 μg/ ml) and showed the lowest cell viability at the highest CSNPs concentration (1000 μg/ ml) (Zaki *et al*., 2015). These findings are not in agreement with the findings of the present study and clearly support the non-cytotoxic effect of the particles on Vero-cell.

## SUMMARY AND CONCLUSION

The *in-vitro* study was conducted using different concentration of nano/microparticle in Vero cell line with maximum final concentration of 1000 μg/ml. The morphological examination showed to maintain the cell monolayer intact with minimal to no loss of cellular integrity and cellular architecture. No significant influences of Gantrez® nanoparticle or chitosan nanoparticles or PLG microparticle were found on alteration of cellular architecture. However, the increasing concentration of PLG microparticle was found to negatively impact on cell viability and cellular metabolic activities (evident from MTT assay). The chitosan nanoparticles and Gantrez^®^ nanoparticles do not appear to have any significant impact on either cell viability or in cellular metabolic activities. Based on the present findings, the conclusions can be drawn as chitosan nanoparticles and Gantrez^®^ nanoparticles do not influence the cell viability and metabolic activities of Vero cell line up to the concentration of 1000 μg/ml. On the other hand, the PLG microparticles have negative impact on cell viability and metabolic activities of Vero cell line.

